# Escherichia-Shigella expansion and metabolite dysregulation in type 3c diabetes

**DOI:** 10.64898/2025.12.02.691786

**Authors:** Erika Kvalem Soto, Valeria Wagner, Julia Engl, Michael Mederer, Veronika Cibulkova, Johanna Piater, Benedikt Schäfer, Elena Dunzendorfer, Silvio Waschina, Susanne Kaser, Konrad Aden, Zlatko Trajanoski, Herbert Tilg, Maria Effenberger

## Abstract

Emerging evidence supports a bidirectional gut–pancreas axis in which microbial dysbiosis, barrier dysfunction, and altered metabolite fluxes contribute to pancreatogenic diabetes (T3cDM). Whether gut microbial changes reflect systemic metabolic disturbances or primarily arise from exocrine pancreatic insufficiency (EPI) remains unclear. We profiled the gut microbiome of 48 outpatients with T3cDM, type 1 diabetes (T1DM), and healthy controls. Genus-level 16S rRNA data were analysed using cross-validated LASSO logistic regression and patient-specific community metabolic models. T3cDM showed reduced α-diversity and distinct β-diversity compared with T1DM and controls. Key compositional shifts included enrichment of *Enterobacteriaceae* (notably *Escherichia–Shigella*) and *Streptococcaceae* in T3cDM. LASSO models discriminated T3cDM from T1DM (AUC 0.867; accuracy 0.818), highlighting *Blautia, Escherichia–Shigella, Streptococcus, Clostridium, and Faecalibacterium* as predictors. Metabolic modelling indicated elevated *Escherichia–Shigella* growth in T3cDM and disease-specific metabolite fluxes. Gut microbial shifts in T3cDM predominantly reflect EPI rather than systemic metabolic disturbances characteristic of T1DM, underscoring the central role of exocrine pancreatic dysfunction in shaping the gut microbiome and its metabolic activity.

## 1. Introduction

Pancreatogenic diabetes mellitus (T3cDM) is defined as diabetes secondary to exocrine pancreatic disease, including total or partial pancreatectomy, pancreatic agenesis, acute and chronic pancreatitis, cystic fibrosis, haemochromatosis, and pancreatic ductal adenocarcinoma [1]. Despite representing only a minor fraction of pancreatic volume, islet cell loss or dysfunction frequently leads to impaired glucose homeostasis in these conditions. Although grouped under a single entity, the pathophysiology varies by etiology and has been best characterised in chronic pancreatitis, pancreatic cancer, and cystic fibrosis [2–4].

The recognition of bidirectional interactions between the gut microbiota and pancreatic diseases has significantly advanced our understanding of pancreatic pathophysiology [5,6]. Emerging evidence indicates that bacterial translocation from the small intestine to the pancreas plays a pivotal role in linking microbial dysbiosis to pancreatic inflammation and malignancy [7,8]. Gut-derived microbes may migrate via the pancreatic duct, lymphatic drainage, or systemic circulation, establishing local pancreatic colonization that perturbs tissue homeostasis and influences disease progression [7,9]. Among these microbial shifts, an overrepresentation of *Escherichia* species has been consistently observed in conditions associated with exocrine pancreatic dysfunction, highlighting that specific taxa may preferentially exploit altered pancreatic and intestinal niches [5,10].

Accumulating data suggest that gut-derived microbes modulate pancreatic disease progression through both local and systemic immune responses [7,9,10]. Microbial components can activate pattern recognition receptors on pancreatic epithelial and immune cells, triggering the release of pro-inflammatory cytokines and chemokines that contribute to tissue injury. They also influence the production of antimicrobial peptides, alter the composition and activity of intrapancreatic immune cell populations, and modulate systemic inflammatory pathways, amplifying pancreatic damage. Notably, the pancreas does not harbor a resident microbiota under normal conditions; thus, dysbiosis in the gut may act as a primary driver of pancreatic inflammation and exocrine pancreatic insufficiency (EPI) [5,9]. These findings underscore the critical role of the gut–pancreas axis in modulating both local injury and broader disease manifestations.

The metabolome has emerged as a key mediator linking pancreatic dysfunction to systemic metabolic consequences. In T3cDM, alterations in lipid, amino acid, and bile acid metabolism reflect both endocrine impairment and EPI [11,12]. Distinct metabolomic signatures have been reported in acute and chronic pancreatitis, with shifts in energy substrates and oxidative stress pathways paralleling disease severity [13,14]. Disrupted amino acid and branched-chain fatty acid profiles in T3cDM suggest combined contributions of β-cell loss, glucagon deficiency, and maldigestion [15]. Additionally, shifts in bile acids and SCFA pools emphasize the contribution of gut–liver–pancreas crosstalk to glucose dysregulation [16]. Current hypotheses propose a bidirectional axis in which microbial metabolites, particularly SCFA, influence pancreatic exocrine and endocrine function, while pancreatic secretions reciprocally shape the composition and activity of the gut microbiota. Pancreatic injury is associated with characteristic microbial shifts—including enrichment of *Escherichia*—barrier dysfunction and bacterial translocation, with the severity of these alterations correlating with both EPI and systemic complications [1,17]. However, it remains unclear whether gut microbial signatures predominantly reflect exocrine pancreatic dysfunction or the metabolic consequences of islet cell loss in T3cDM.

Here, we compared T3cDM and T1DM longitudinally to identify microbial community features unique to T3cDM. Using in silico modelling and machine learning models, we predicted metabolic exchanges within the gut microbiota and linked these to clinical outcomes. We observed a pronounced enrichment of *Escherichia–Shigella* in T3cDM, correlating with pancreatic functional loss and EPI rather than the metabolic changes seen in T1DM, alongside distinct microbial lipid and carbohydrate production profiles. These findings support a model in which the gut microbiome functions as a dynamic metabolite pool that senses and modulates pancreatic dysfunction.

## 2. Results

### 2.1 Reduced diversity and distinct community structures in T3cDM compared with T1DM and healthy controls

Clinical characteristics of the cohorts are provided in Supplementary tables 1, 2 and 3. Microbial diversity was significantly reduced in T3cDM compared with both T1DM and healthy controls, as shown across multiple α-diversity indices. The Simpson diversity index differed between healthy controls and T3cDM (p.adj < 0.05), and Shannon entropy differed between T1DM and T3cDM (p.adj < 0.05) (Figure 1A). β-diversity analyses supported these observations. Bray–Curtis dissimilarity revealed separation between T1DM and T3cDM, with healthy controls clustering in between (Figure 1B.1). Jaccard-based PCoA provided clearer separation of T1DM and T3cDM, while T1DM partially overlapped with healthy controls (Figure 1B.2). In total, 132 microbial families were detected, with 86 shared across all groups (Figure 1C). Twenty families were unique to T1DM and five to T3cDM. Principal component analysis showed greater dispersion in T1DM and T3cDM compared with controls, with pronounced heterogeneity in T1DM (Figure 1D).

**Figure 1.**
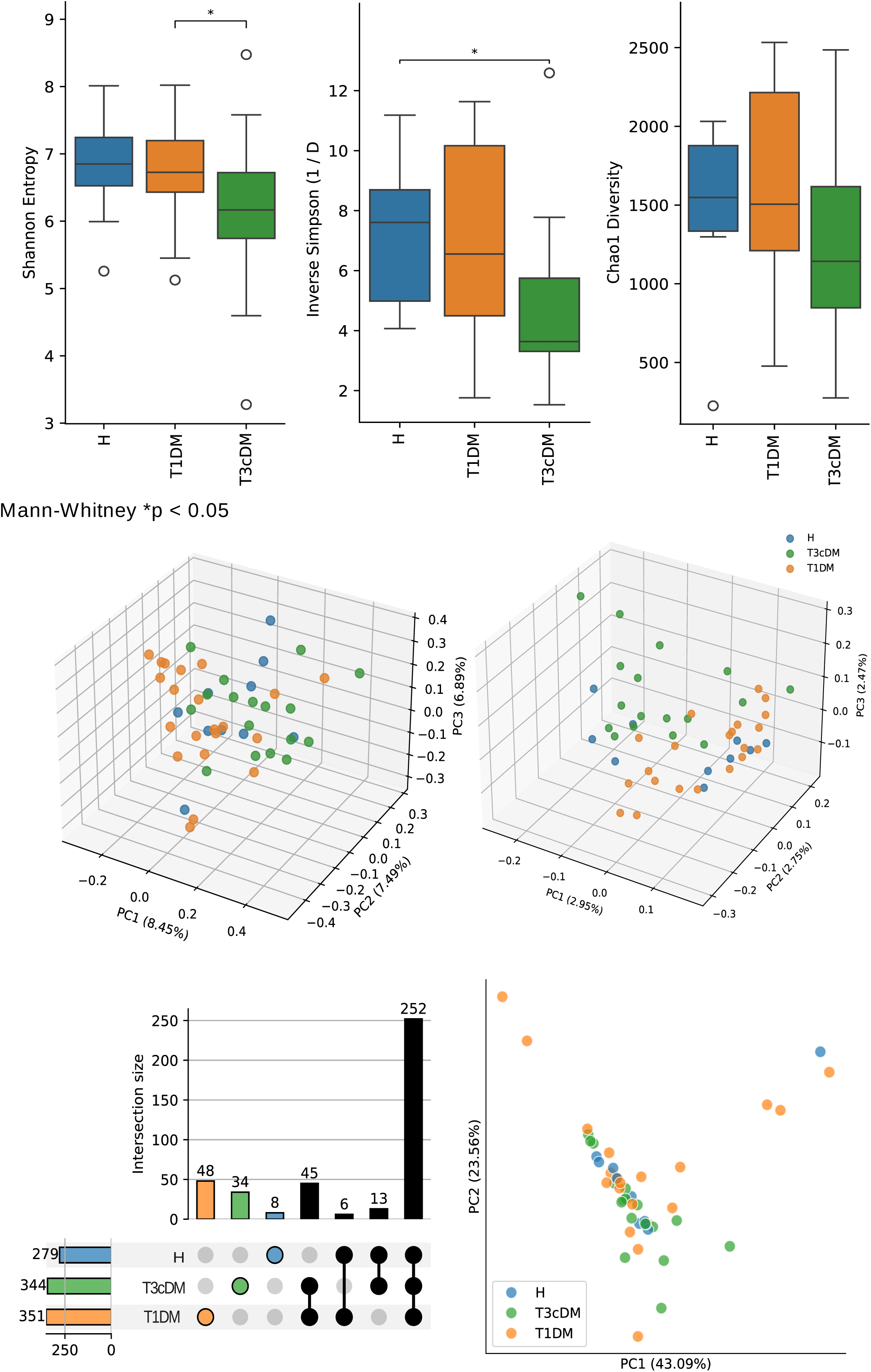
Bacterial taxa diversity and family overlap. (A)Community α-diversity analysis described by Shannon, Simpson and Chao1 index measurements, statistical annotation indicates p.adj < 0.05 after two-sided Mann Whitney test and FDR correction. (B) Community Ꞵ-diversity analysis described by Bray-Curtis and Jaccard distances. All analyses are separated by condition: Healthy (H), T1DM (Type 1 Diabetes Mellitus), T3cDM (Type 3C Diabetes Mellitus). (C) UpSetPlot visualizing the set overlap for all the recorded families. (D) Principal component analysis (PCA) for all the samples where the two principal components account for most of the variance.

### 2.2 Disease-specific shifts at the family level

The ten most abundant families in T3cDM included *Bifidobacteriaceae, Bacteroidaceae, Barnesiellaceae, Prevotellaceae, Rikenellaceae, Lachnospiraceae, Oscillospiraceae, Ruminococcaceae, Enterobacteriaceae*, and *Akkermansiaceae* (Figure 2A), consistent with taxonomy assignments shown in the Sankey diagram (Figure S1). Core microbiome analysis identified eight families consistently shared across groups (Figure 2B). Pairwise comparisons of these core families revealed disease-specific differences (Figure 2C). *Enterobacteriaceae* and *Streptococcaceae* were significantly enriched in T3cDM compared with both T1DM and healthy controls. *Bacteroidaceae* were enriched in T1DM compared with the other groups, whereas *Ruminococcaceae* were enriched in healthy controls.

**Figure 2.**
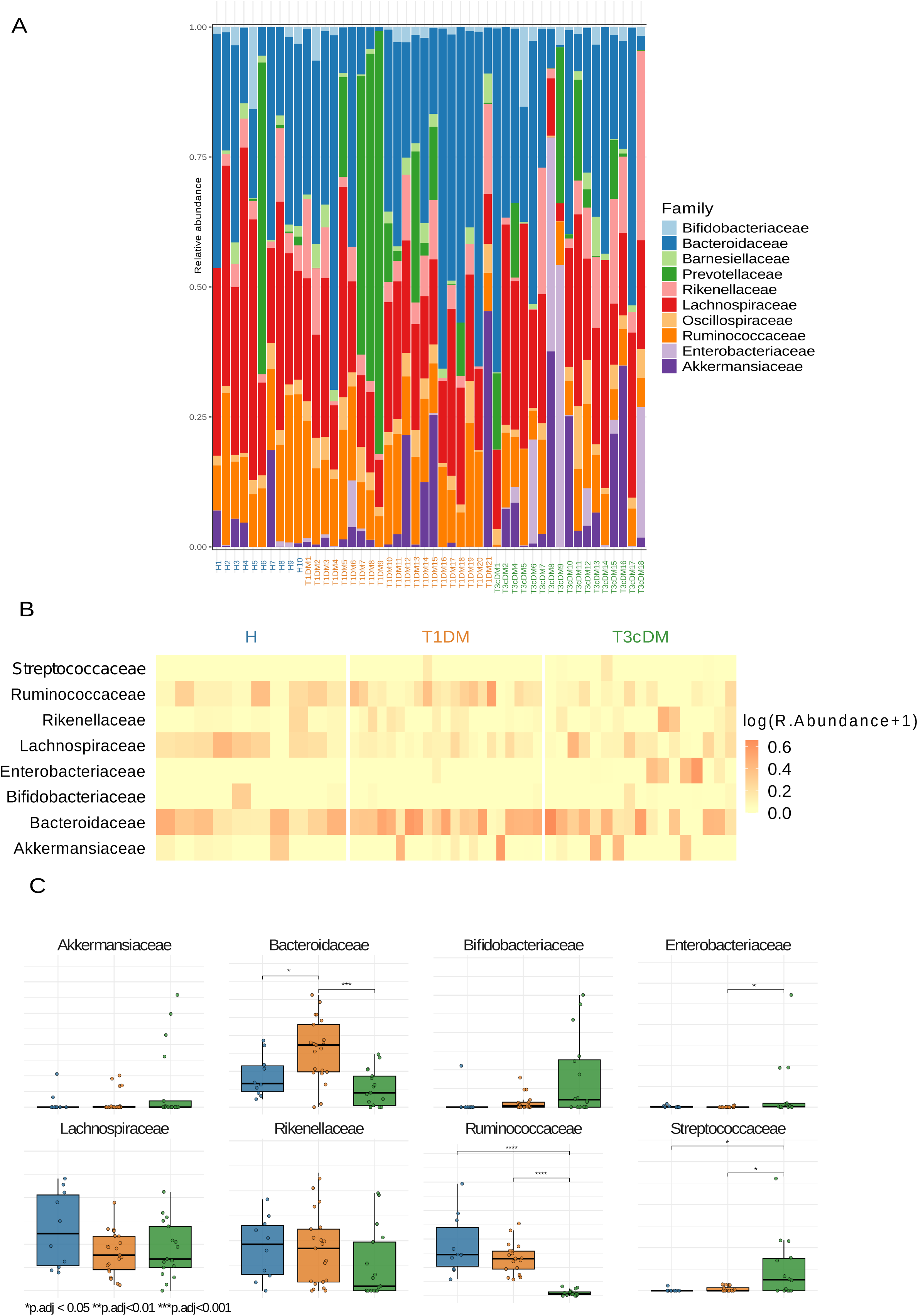
Family-level bacterial composition. (A) The stacked bar plot depicts the relative abundance for the 10 most abundant families in all samples. (B) Heatmap showing the log(relative abundance +1) for the core microbiome, defined as the bacterial families present in more than 50% of the samples. (C) Boxplots for the log(relative abundance +1) of the core microbiome families together with pairwise comparison across the disease groups using Mann–Whitney test statistical annotation*p.adj < 0.05, ** p.adj < 0.01, *** p.adj < 0.001.

### 2.3 Logistic regression and LEfSe identify microbial signatures of T3cDM

To identify discriminatory taxa, logistic regression was applied at the genus level. For T1DM vs. T3cDM (primary comparison), the confusion matrix showed robust classification with a ROC AUC of 0.867 and accuracy of 0.818 in the test set (Figure 3A–B). The most influential genera included *Blautia, Escherichia–Shigella, Streptococcus, Clostridium*, and *Faecalibacterium* (Figure 3C). Overlap across models was visualized by Venn diagram (Figure 3D), and relative abundances confirmed that *Blautia, Faecalibacterium,* and *Subdoligranulum* were depleted in both diabetes groups, whereas *Streptococcus* and *Escherichia–Shigella* were enriched in T3cDM (Figure 3E). LEfSe analysis confirmed discriminatory genera across pairwise comparisons (Figure S2), overlapping with taxa identified by regression models. *Faecalibacterium prausnitzii* and *Subdoligranulum* were more abundant in healthy controls than in T1DM, whereas *Streptococcus* and *Escherichia–Shigella* were enriched in T3cDM. Sex-specific analyses further revealed distinct microbial signatures (Figure S3): *Turicibacter* was enriched in T1DM males, *Blautia* in T1DM females. In T3cDM, *Ruminococcus* and *Subdoligranulum* were enriched in males while *Eggerthella*, *Eubacterium*, and *Negativibacillus* were enriched in females.

**Figure 3.**
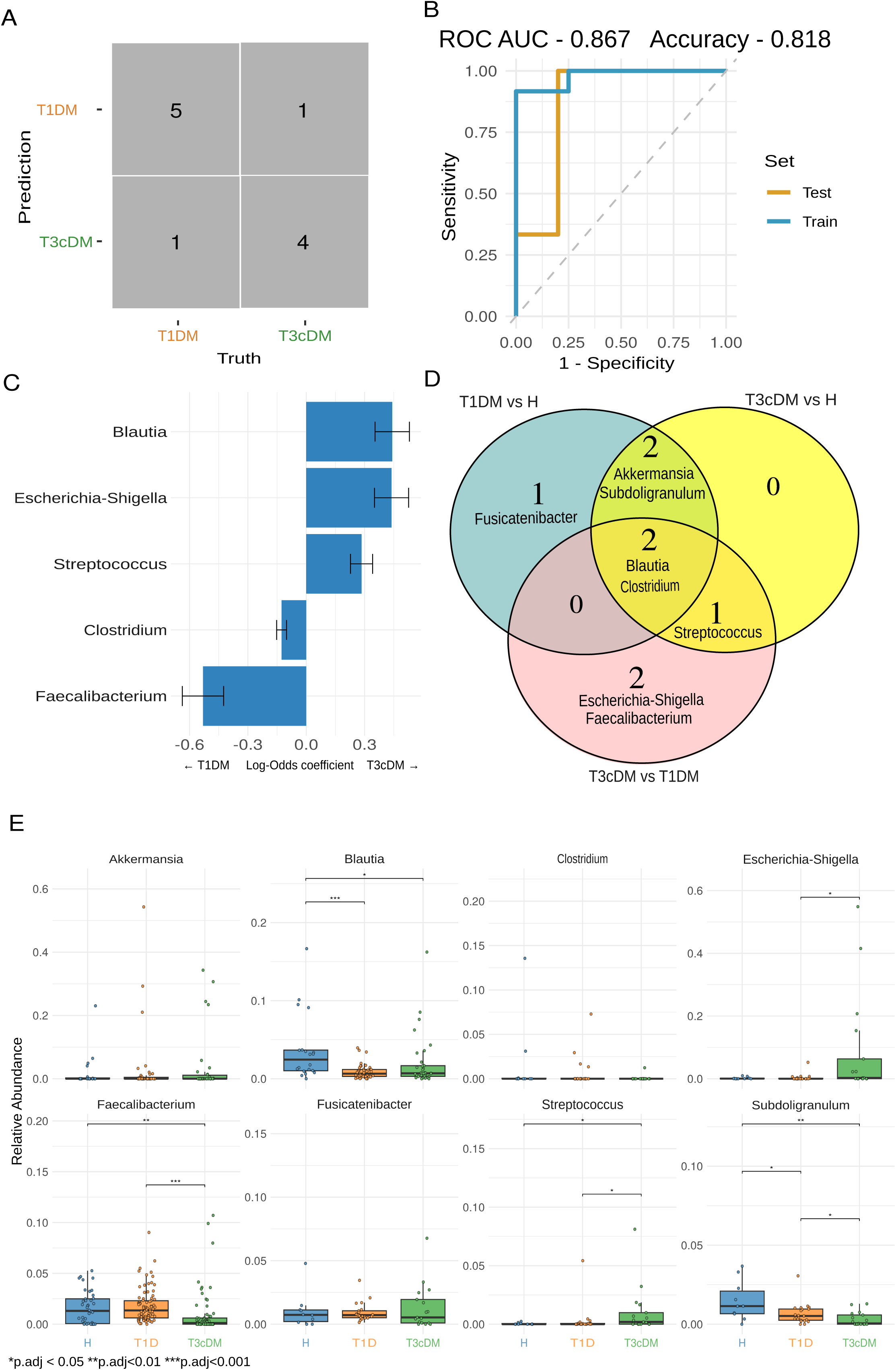
Microbial signatures identified by logistic regression. (A) Confusion matrix evaluating the performance of the microbial logistic regression model for the classification between T1DM and T3cDM on the test set. (B) Receiver operating characteristic (ROC) curve showing the performance of the classification model. Visualizes the relationship between the true positive rate (sensitivity) and the false positive rate (1 - specificity). The model achieved an area under the ROC curve (AUC) of 0.867 and an accuracy of 0.818 on the test set. (C) Bar plot showing the top five genera contributing to the classification model based on their log-odds coefficient. Positive coefficients indicate genera associated with increased odds of classification into the T3cDM, while negative coefficients suggest association with T1DM. (D) Venn diagram showing the overlap of the top five genera from pairwise classification models comparing T1DM vs. T3cDM, T1DM vs. H, and T3cDM vs. H. (E) Boxplots plots showing relative abundance of microbial genera identified as top contributors in the logistic regression model from all the pairwise classification models.

Further analyses included model validation across all pairwise comparisons (Supplementary material). Confusion matrices demonstrated good classification performance for T1DM vs. controls and T3cDM vs. controls (Figure S4). ROC curves yielded AUCs of 0.867 (accuracy 0.75) for T1DM vs. controls, and 0.833 (accuracy 0.70) for T3cDM vs. controls (Figure S5). The top five genera contributing to classification were visualized for both comparisons (Figure S6).

Correlation analyses highlighted strong clinical associations. *Escherichia–Shigella* abundance correlated closely with pancreatectomy (Figure S7). Its abundance was significantly higher in individuals with pancreatectomy, but did not differ between left, right, or total resections (Figure S8). Furthermore, *Escherichia–Shigella* abundance was significantly correlated with stool elastase, the most reliable marker of EPI (Figure S9).

Longitudinal clinical changes were evaluated by generalized linear mixed-effects models (Figure S10). In T1DM, NT-proBNP, alkaline phosphatase, high-sensitivity troponin T, and body-mass index (BMI) increased over a 5-year period, whereas in T3cDM, LDL cholesterol decreased modestly. Furthermore, BMI did not differ significantly between groups (p = 0.896), but metabolic dysfunction-associated steatotic liver disease (MASLD), coronary artery disease (CAD), and arterial hypertension were significantly more prevalent in T3cDM (p = 0.015, p = 0.014, and p < 0.001, respectively). Use of antihypertensive agents and diuretics was also more frequent in T3cDM (p < 0.001 and p = 0.019). Detailed characteristics are provided in Supplementary tables 1, 2 and 3.

### 2.4 Functional modeling reveals altered growth dynamics and metabolite flux in T3cDM

Community metabolic modelling revealed functional perturbations underlying compositional shifts. Flux-based clustering further resolved microbial functional differences (Figure S11). *Akkermansia* and *Blautia* were predominantly associated with T1DM, *Escherichia–Shigella* with T3cDM, whereas *Faecalibacterium*, *Streptococcus*, and *Subdoligranulum* showed more variable distributions. Predicted genus-specific growth rates from community metabolic simulations are shown for the 20 most abundant genera (Figure S12) and indicated clear genus-level differences (Figure 4A; Figure S12). *Escherichia–Shigella* had the highest growth rate in T3cDM and the lowest in healthy controls. *Blautia* growth was reduced in T1DM, and *Faecalibacterium prausnitzii* growth was lowest in T3cDM. Predicted metabolite abundances revealed enrichment of nucleotides (cytosine, deoxyguanosine, nicotinamide mononucleotide), carbohydrates (starch, pectins), hydrogen, and SCFA (propionate, butyrate) in T3cDM (Figure 4B). In contrast, several carbohydrates (D-fructose, D-mannose, N-acetylneuraminate), amino acid–related metabolites (putrescine, glutathione), lipids, carboxylic acids, bile acids, and SCFA (acetate, formate) were depleted. In more detail, we can see connected genera of interest to metabolite predicted abundance (Figure S13). *Akkermansia*, *Blautia*, *Faecalibacterium*, and *Subdoligranulum* are connected to 2-oxobutanoate, D-galactose, N-acetylneuraminate, sodium, cellobiose, D-fructose, N-acetyl-D-glucosamine, D-glucuronate, glycerol, acetaldehyde, butyrate, acetate, ammonium, and calcium. *Escherichia–Shigella* and *Streptococcus* are linked to formate, 2-oxoglutarate, choline, trehalose, adenosine, chloride, copper, and CO₂. Linear regression further identified *Escherichia–Shigella* as uniquely associated with a broad metabolite spectrum including inorganic ions, nucleotides, amino acid–related metabolites, carbohydrates, carboxylic acids, and vitamins (Figure 4C). Together, these findings demonstrate that functional modeling resolves disease-specific metabolic alterations not evident from taxonomic profiles alone.

**Figure 4.**
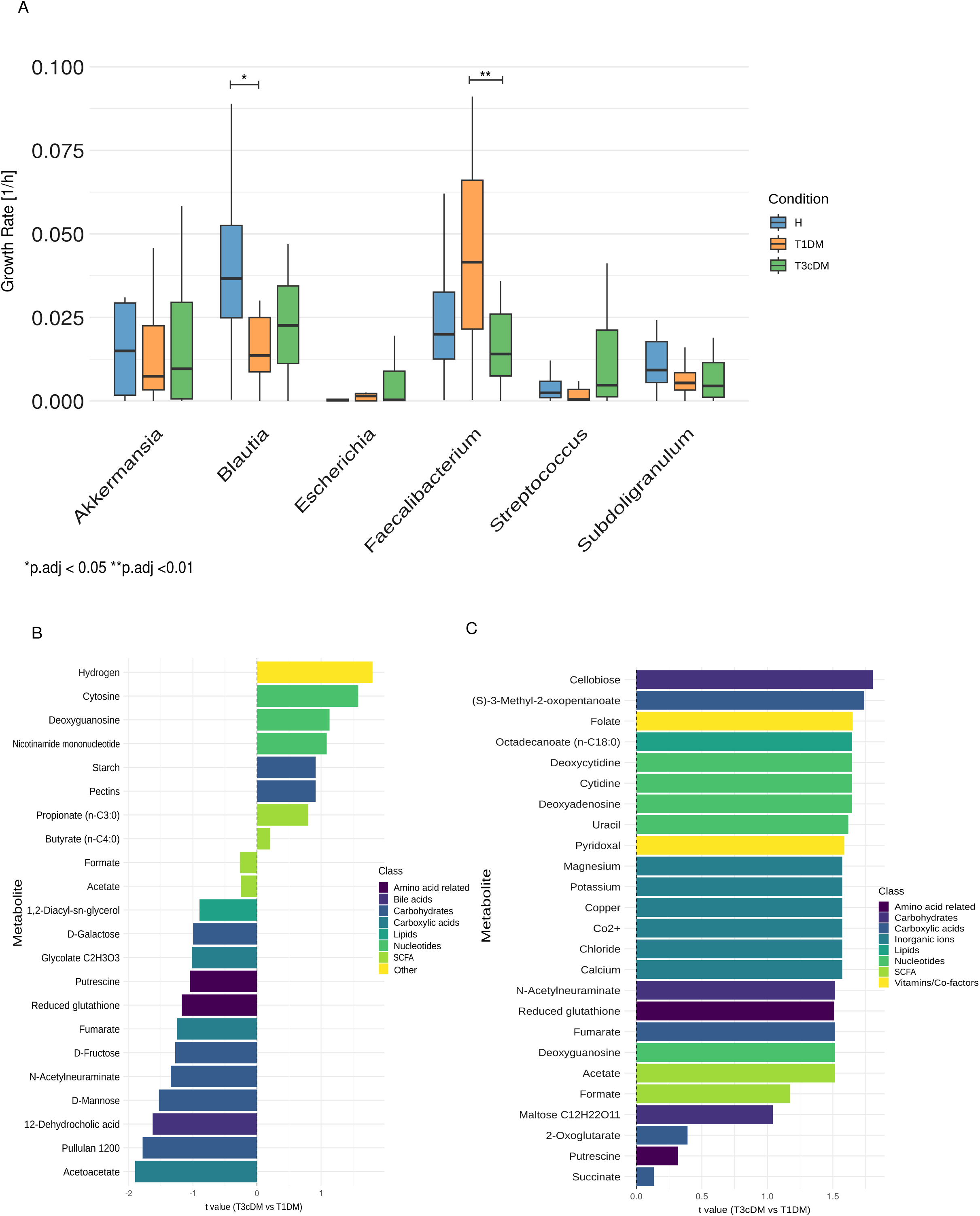
Microbial community model metabolic in-silico prediction. (A) Boxplots showing the predicted growth rates of bacterial genera previously identified by logistic regression. (B) Barplots showing linear regressing analysis of metabolites abundance for all identified genera and exclusively for Escherichia-Shigella (C) between T3cDM vs T1DM, adjusted for sex as confounder. Positive t value indicated higher concentrations in T3cDM and color code indicates metabolite class. Significance was defined as *p*.adj < 0.05 (Wilcoxon test).

## 3. Discussion

There is broad consensus that both endocrine and exocrine pancreatic insufficiency are linked to intestinal dysbiosis, manifesting as altered microbial composition and reduced α-diversity. However, causal relationships in humans remain elusive, and the downstream effects of dysbiosis on host immunity are only partially understood. Although microbial perturbations have been described in chronic pancreatitis, post-pancreatectomy states, and T1DM, the gut microbiome in T3cDM—in which endocrine failure is secondary to exocrine loss—has been insufficiently characterized to date [24–26].

Consistent with prior work, we observed pronounced dysbiosis in T3cDM, characterized by a marked loss of microbial diversity and selective enrichment of facultative pathobionts such as *Enterobacteriaceae* and *Streptococcaceae* [24–28]. Similar microbial shifts have been reported in chronic pancreatitis, pancreatic cancer, and EPI, conditions in which impaired digestion and nutrient malabsorption favor expansion of opportunistic taxa and small intestinal bacterial overgrowth (SIBO) [24,27,31]. In contrast, the gut microbiota in T1DM more closely resembles healthy controls, showing preserved α-diversity and enrichment of immune-associated taxa rather than a broad community collapse [29]. The markedly lower diversity observed in T3cDM likely reflects the combination of long-standing disease, chronic inflammation, maldigestive conditions, and progressive exocrine loss [30,31].

SCFA–producing genera such as *Blautia*, *Faecalibacterium*, and *Subdoligranulum* were consistently depleted in T3cDM, while facultative pathobionts including *Streptococcus* and *Escherichia–Shigella* were selectively enriched [27,28,32]. Expansion of *Enterobacteriaceae* coincided with reduced predicted growth of *Blautia* and *Faecalibacterium*, suggesting loss of protective microbial functions. *Escherichia–Shigella* reached its highest relative abundance in patients with low stool elastase, implicating EPI as a key ecological driver. Experimental evidence supports this observation: enzyme deficiency promotes *Enterobacteriaceae* expansion, which can be mitigated by pancreatic enzyme replacement therapy (PERT) [30.42]. These findings collectively indicate that the exocrine defect in T3cDM acts as a dominant selective force reshaping microbial ecology.

Metabolomic profiling revealed a distinct metabolic signature in T3cDM, characterized by reduced metabolite diversity, a dominance of hydrogen and nucleotide pathways, and limited SCFA and carbohydrate contributions. By contrast, T1DM samples displayed higher representation of carbohydrate, amino acid, and carboxylic acid pathways but similarly showed diminished SCFA levels, consistent with altered fermentation in autoimmune diabetes [29]. Nucleotide enrichment in T3cDM suggests accelerated microbial turnover, while elevated polysaccharide residues likely reflect impaired enzymatic hydrolysis in the context of EPI [27,31]. Hydrogen production, driven by fermentation of undigested carbohydrates, was a dominant feature, with *Escherichia–Shigella* contributing metabolites such as formate and acetaldehyde that can compromise epithelial barrier integrity [33]. The loss of butyrate-producing taxa further deprives the host of key microbial metabolites essential for maintaining epithelial and immune homeostasis [32,37].

Moreover, enhanced microbial metabolism of choline and glycine betaine may exacerbate cardiometabolic risk through the generation of trimethylamine (TMA) and its hepatic oxidation product, trimethylamine-N-oxide (TMAO) [34–36]. These microbial–host co-metabolites are well-established mediators of atherogenesis and metabolic dysfunction, and their enrichment in T3cDM supports a mechanistic link between exocrine loss, dysbiosis, and systemic metabolic derangements. Although our cohort size was limited, we implemented supervised machine learning models to explore genus-level and metabolic features distinguishing T3cDM, T1DM, and healthy individuals. Recent studies have applied similar approaches to identify microbiome-derived signatures across pancreatic and metabolic diseases. Tan et al. and Yuan et al. integrated gut microbiota, serum metabolites, and lipidomic data in T1DM, revealing multi-omic biomarkers predictive of disease progression [29,41]. Lee et al. and other groups have used machine learning to define microbiome-based diagnostic classifiers in pancreatic cancer and chronic pancreatitis, underscoring the diagnostic potential of AI-driven models [38–40]. While these tools hold significant promise, it is essential to apply rigorous cross-validation, avoid data leakage, and interpret model features biologically rather than solely statistically[39,43]. Our results, in line with these studies, highlight the emerging role of artificial intelligence in delineating microbiome–host interactions and identifying mechanistic pathways in metabolic and pancreatic diseases.

The dysbiotic profile observed in T3cDM—marked depletion of SCFA producers and enrichment of facultative pathobionts—likely contributes to mucosal barrier disruption and systemic inflammation. SCFA such as butyrate and propionate regulate epithelial integrity, maintain regulatory T cell homeostasis, and modulate inflammatory signaling via HDAC inhibition and NF-κB suppression [32,37]. Loss of these microbial functions may permit translocation of bacterial products, triggering TLR and NLRP3-mediated immune activation and metabolic inflammation. Meanwhile, *Escherichia–Shigella*–derived metabolites such as acetaldehyde and formate can further compromise epithelial tight junctions and induce oxidative stress [33]. Together, these alterations may promote chronic low-grade inflammation and metabolic dysfunction typical of T3cDM.

Our study provides an integrative microbiome–metabolome characterization of T3cDM, delineating a distinct dysbiotic and metabolic profile that primarily reflects exocrine insufficiency rather than isolated endocrine dysfunction. The enrichment of facultative pathobionts, particularly *Escherichia–Shigella* and *Streptococcaceae*, together with the depletion of SCFA-producing genera such as *Blautia* and *Faecalibacterium*, defines a microbial ecosystem conducive to metabolic dysregulation, epithelial barrier disruption, and systemic inflammation. However, the cross-sectional nature of the analysis precludes causal inference. Longitudinal and interventional studies—especially those incorporating pancreatic enzyme replacement therapy, dietary modulation, or targeted microbiome interventions—are needed to clarify causal relationships between dysbiosis, exocrine dysfunction, and metabolic outcomes. Future integrative approaches combining metagenomic, transcriptomic, and immune-phenotyping data will be essential to identify strain-level microbial effectors driving host–microbe interactions. In parallel, robust artificial intelligence frameworks with external validation should be employed to enhance biomarker discovery, refine predictive modeling, and advance microbiome- and metabolome-informed diagnostics and therapeutics in T3cDM and related disorders [38–40,43].

## 4. Methods

### 4.1 Study population

We enrolled 48 participants: 21 with T1DM, 17with T3cDM, and 10 healthy controls with a follow up.

Definitions of T3cDM and T1DM followed previously published consensus criteria.^18,19^ Healthy controls were selected based on the absence of comorbidities and no regular medication use. The T1DM and T3cDM group were 100% insulin dependent. The mean interval between baseline clinical and laboratory assessment and the second follow-up was 2411.1 (SD ± 323.8) days, and stool samples were obtained a mean of 571.2 (SD ± 339.0) days after baseline. Sex distribution did not differ significantly between groups. T1DM patients were significantly younger than both T3cDM and healthy participants, whereas smoking and daily alcohol intake > 20 g were more common in T3cDM. Within the T3cDM group, 13.0% (n = 3) had undergone total pancreatectomy, 39.1% (n = 9) partial pancreatectomy (pancreatic head or distal resection), and 47.8% (n = 5) had chronic pancreatitis.

### 4.2 Machine learning modelling

We trained LASSO (glmnet, mixture=1) logistic regression models using the tidymodels framework in R (v4.3.0) on genus-level microbial profiles derived from the nf-core/ampliseq pipeline.^20,21^ Taxa were aggregated at the genus level, filtered (mean relative abundance >3%, prevalence >5%), and normalised. Samples were annotated into three groups (T1DM, T3cDM, healthy) based on metadata. Separate binary classification models were trained for each comparison, with cross-validation applied. Model performance was evaluated by accuracy, AUC (area under the curve), and ROC (receiving operating curve) curves, while feature importance was inferred from non-zero model coefficients.

### 4.3 Microbial community modelling

Community metabolic modelling (CMM) was employed to integrate genome-scale metabolic models of individual microbial species, enabling the simulation and prediction of interspecies metabolic interactions, nutrient exchanges, and overall community function within a shared metabolic environment. CMM was performed with MICOM v0.25.1 [22]. Taxonomic profiles from nf-core/ampliseq outputs were collapsed to the genus level, filtering for relative abundance ≥2.5%. Community models were constructed with AGORA v1.03 genus-level metabolic models [23]. A standardised Western diet was applied to simulate microbial growth and metabolite exchange using parsimonious flux balance analysis (pFBA). Sample specific growth rates and exchange fluxes were computed. Growth rate differences between groups were assessed using pairwise t-test with Benjamini–Hochberg correction, and effect sizes calculated. Metabolites identified by logistic regression were plotted as boxplots; statistical significance was defined as FDR-adjusted p<0.05. Linear regression models adjusted for sex were applied to test differences in metabolite abundances between T1DM and T3cDM, with positive t-values indicating higher levels in T3cDM. To explore clustering of metabolic variation by taxon or disease, t-distributed stochastic neighbour embedding (t-SNE) was applied to reduced flux profiles.

### 4.4. Ethical considerations

All research was conducted in accordance with both the Declarations of Helsinki and Istanbul. The institutional ethics commission (Ethics Commission of the Medical University of Innsbruck) approved the study protocol AN 1100/2025 and 1156/2018, written consent was given by all subjects.

### 4.5. Data availability

Microbiome sequencing data and statistical analysis results are available in Zenodo (https://doi.org/10.5281/zenodo.16794434). All scripts used for data analysis, statistics, and visualizations are provided through the GitHub repository (https://github.com/icbi-lab/diabetes_microbe).

More detailed information for material and methods is available in the supplementary data.

## 5. Author contributions

ME designed the project. VW and EKS wrote the paper. EKS verified the analytical methods and analyzed the data. ED contributed to clinical data acquisition. SW and ZT contributed to the analytical methods. VW, EKS and ME contributed to the interpretation of the results. JE contributed to patient recruitment sample preparation. VW, EKS, ME, HT, TA, BS, JE, SW, ZT, JP, VC, KA, MM and SK provided critical feedback and contributed to data analysis and manuscript preparation.

## 6. Acknowledgement

We thank Ursula Bachlechner for her assistance in acquiring patient stool samples and performing all subsequent laboratory processing. Further we are grateful for her assistance in assessing clinical parameter of the study cohort.

The author used OpenAI’s ChatGPT (OpenAI, GPT-5, 2025) to assist in improving the language style and clarity of the English text. The model was used for grammar correction, professional phrasing, and enhancing the overall readability. All scientific content, ideas, and interpretations are solely the author’s own. Prompt example used: *“Please revise the following paragraph to sound more professional and fluent in academic English:“*

## 7. Conflict of interest

None declared.

## 8. Funding

This work is supported by the excellence initiative VASCage (Centre for Promoting Vascular Health in the Ageing Community), an R&D K-Centre (COMET program—Competence Centers for Excellent Technologies) funded by the Austrian Ministry for Transport, Innovation and Technology, the Austrian Ministry for Digital and Economic Affairs, and the federal states Tyrol, Salzburg, and Vienna. ME was supported by the Austrian Society of Gastroenterology and Hepatology (Science Funding 2022) and the Land Tirol (KI in Gesundheitswesen). VW was gratefully supported by an award from the Austrian Society of Gastroenterology and Hepatology (Dr. Falk Förderung für Jungwissenschaftler, ÖGGH). JE was gratefully supported by the MUI Start program funded by the Medical University of Innsbruck. EKS was supported by the PhD programs Molecular and Cellular Basis of Diseases (MCBD) and CONNECT from the Medical University of Innsbruck.

*No funders were involved in the study concept, design, conduct, data analyses, writing of the manuscript, or the decision to submit this work for publication*.

## 10. Glossary

ASVs: Amplicon sequence variants
ANOVA: Analysis of variance
BMI: Body mass index
CAD: Coronary artery disease
EPI: Exocrine pancreatic insufficiency
Iron: Fe³⁺
GLMM: Generalized linear mixed-effects models
GM: Gut microbiome
H: Healthy control
IBD: Inflammatory bowel diseases
IQR: Inter-quartile range
LEfSe: Linear discriminant analysis Effect Size
MASLD: Metabolic dysfunction-associated steatotic liver disease
NODAP: New-onset diabetes after pancreatitis
PCR: Polymerase chain reaction
T3cDM: Pancreoprivic diabetes
ROC: Receiver operating characteristic
SCFA: Short-chain fatty acids
SD: Standard deviation
TMA: Trimethylamine
TMAO: Trimethylamine N-oxide
T1DM: Type 1 Diabetes

## Appendices

## A. Research in Context

### Evidence before this study

We searched PubMed up to September 2025 for studies examining the gut microbiome in pancreatogenic diabetes (T3cDM). While numerous studies have reported microbial dysbiosis in chronic pancreatitis, pancreatic cancer, post-pancreatectomy patients, and type 1 diabetes (T1DM), data in T3cDM are limited. Most prior work focused on cross-sectional associations between microbial composition and metabolic parameters, with few studies integrating functional metabolomics or mechanistic modelling.

### Added value of this study

Our study provides a comprehensive, longitudinal comparison of gut microbiota and microbial metabolic function in T3cDM, T1DM, and healthy controls. Using community metabolic modelling, we identify a distinct microbial and metabolomic signature in T3cDM, characterised by enrichment of *Escherichia–Shigella* and Streptococcaceae, depletion of short-chain fatty acid (SCFA)-producing genera, and metabolic shifts such as increased hydrogen and nucleotide production. These alterations correlate with exocrine pancreatic insufficiency rather than endocrine dysfunction, highlighting EPI as a key driver of gut microbial dysbiosis and metabolite perturbation.

Our findings support a mechanistic gut–pancreas axis in T3cDM, linking EPI to microbial and metabolic changes that may contribute to chronic inflammation and cardiometabolic risk. Integration of microbial and metabolomic data offers potential for novel diagnostic biomarkers and therapeutic strategies targeting the microbiome to mitigate complications in T3cDM.

